# Why they move: breeding-site availability drives partial migration in a large terrestrial reptile

**DOI:** 10.64898/2026.08.20.745942

**Authors:** Marco Gargano, Lorenzo Garizio, Giuliano Colosimo, Pierpaolo Loreti, Alexandro Catini, Lorenzo Bracciale, Massimiliano De Luca, Gregory A. Lewbart, Christian Sevilla, Glenn P. Gerber, Paolo Gratton, Gabriele Gentile

## Abstract

1. Migration is a widespread phenomenon across taxa, yet the ecological mechanisms underlying its evolution and maintenance, particularly whether migratory behaviors are primarily driven by access to spatially restricted breeding sites or by seasonal tracking of trophic resources, remain poorly documented outside birds and large mammals. Despite increasing evidence that reptiles perform seasonal migrations, the ecological mechanisms underlying these movements have rarely been formally tested.
2. The critically endangered Galápagos pink land iguana (*Conolophus marthae*), endemic to Wolf Volcano, Isabela Island, exhibits partial migration along a steep altitudinal gradient, providing an opportunity to disentangle the relative roles of breeding-site availability, trophic resource dynamics, and thermoregulatory conditions as drivers of migration.
3. We used GPS tracking data from 22 individuals (7 males, 15 females) monitored between 2019 and 2023, combined with high-resolution spatio-temporal models of vegetation productivity and air temperature across the species’ altitudinal range, to characterize population-level movement patterns and evaluate competing hypotheses explaining the evolution of this migratory behavior.
4. Movement models revealed a clear pattern of partial migration: 16 out of 22 tracked individuals performed seasonal altitudinal movements between a restricted high-elevation mating area and a larger dispersal area at lower elevation, with males reaching the mating area approximately 48 days earlier than females. The dispersal area remained consistently more productive than the mating area throughout the year, rejecting the prediction that individuals should track the shifting trophic resource peaks. Instead, the mating season coincided with the local productivity peak within the mating area, whereas temperature differences between areas were small (*ca.* 2°C) and did not explain migration timing.
5. These results support a site-dependent hypothesis of partial migration over a resource-tracking hypothesis, indicating that access to spatially restricted breeding sites is the primary driver of migration in this species, with local trophic resource dynamics fine-tuning reproductive timing. Providing empirical evidence for the ecological mechanisms underlying migration in a large terrestrial reptile, our results extend site-dependent theories of migration beyond birds and mammals and identify breeding-site availability as a key ecological driver of migratory behaviors across taxa.

## Introduction

Migration is a widespread ecological phenomenon found in a broad range of taxa (Lapin et al., 2025), which influences individual fitness by, among other mechanisms, regulating intra and interspecific competition (Bohrer et al., 2014; Rolandsen et al., 2017; Senner et al., 2019). Although migration has long fascinated researchers, its evolutionary origins across different lineages remain debated (Griswold et al., 2010). One reason is that migratory patterns are still poorly documented for many taxa. Most mechanistic insights into migration are based on studies on birds and large mammals (Soriano-Redondo et al., 2020), while reptiles, except for sea turtles, are still largely understudied (Southwood & Avens, 2010). Expanding our understanding of migration across different taxa is essential to identify the multiplicity of ecological and evolutionary mechanisms underlying this behavior. This need becomes even more urgent in the context of the current anthropogenic pressures, as habitat fragmentation and climate change alter the spatial and temporal synchronization of migration and resources availability, potentially affecting species survival (Bolger et al., 2008; Bohrer et al., 2014; Shaw, 2016).

Migratory behavior is highly variable both among and within species (Dingle et al., 2007; Abraham et al., 2022). While some species and populations perform obligate seasonal movements involving essentially all individuals, others exhibit partial migration, by which some individuals migrate, while others remain resident (Chapman et al., 2011). Partial migration has been documented in many species across diverse taxa, including fish, birds and mammals (Arnekleiv et al., 2022; Berg et al., 2019; Lisi et al., 2022; Martin et al., 2022). From an evolutionary perspective, partial migration has historically been regarded as a preliminary stage in the evolution of obligate migration (Berthold, 1996), providing a powerful system to investigate the ecological and evolutionary drivers of migratory behaviors (Chapman et al., 2011). Theoretical models (Lundberg, 1987; Lundberg, 1988; Kaitala et al., 1993) suggest that partial migration can evolve when individuals face a trade-off between reproductive success and survival, such that the resident strategy promotes higher reproductive success at the breeding site while migratory behavior improves survival during non-breeding season. This would be particularly true when reproductive success relies on the limited presence of suitable breeding areas (Winger et al., 2019), which consequently act as an independent resource, decoupled from foraging opportunities. Salmonid fish are paradigmatic examples of this scenario, because access to suitable spawning habitats strongly constrains reproductive success in this system independently from trophic resource availability (Jonsson & Jonsson, 1993). Under these conditions, frequency-dependent selection can maintain both strategies within the same population, with migrant and resident individuals sharing the same breeding grounds but exploiting different non-breeding habitats (Lundberg, 1987; Lundberg, 1988; Kaitala et al., 1993). This led to the idea that migration is more likely to evolve when access to breeding sites represents the ultimate limiting resource (Kokko & Lundberg, 2001). Under this site-dependent mechanism, it can be expected that migratory behavior may often emerge in animals for which breeding (mating and/or nesting) sites constitute a spatially-clustered limited resource. In such animals, seasonal trophic resource dynamics would act as a secondary layer of selection, fine-tuning migratory and reproductive phenology once the spatial dimension of breeding has been established (Figure 1).

**Figure 1.**
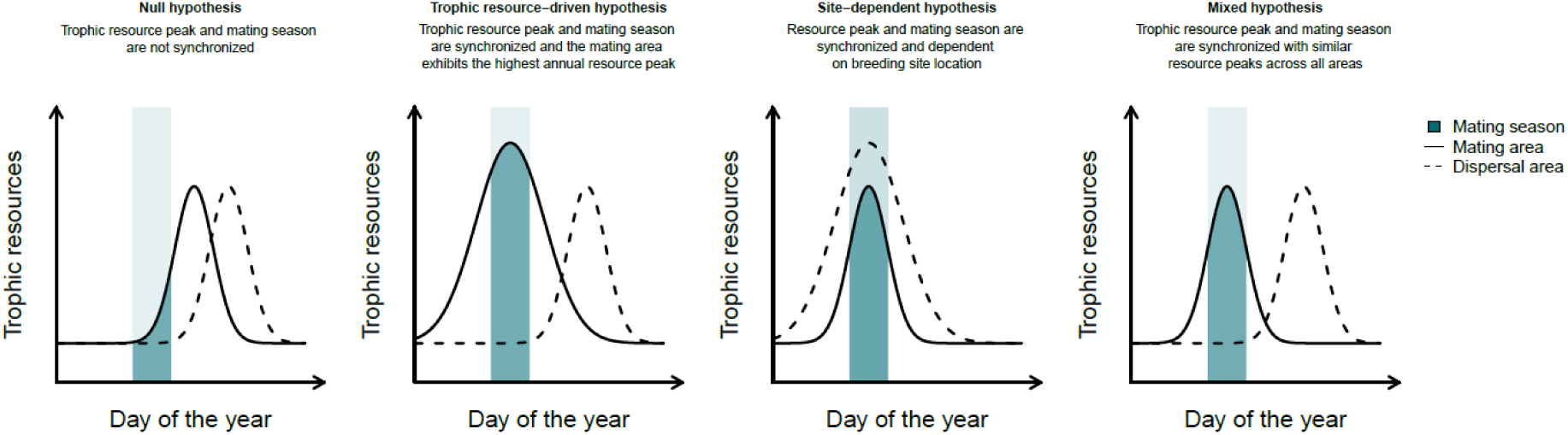
Conceptual expectations linking seasonal trophic resource dynamics and the timing of reproduction under competing hypotheses identifying alternative mechanisms as predominant drivers of migration. The figure illustrates four scenarios describing how the timing of the mating season may relate to seasonal peaks in resource availability across species range. The solid line represents seasonal resource availability in the mating area, whereas the dashed line represents resource availability in the dispersal area. The green shaded vertical band indicates the mating season. Under the null hypothesis, the mating season does not coincide with peak in resource availability in either area. Under the resource-driven hypothesis, migration evolves as a strategy to track spatio-temporal peaks in resource availability. In this case, reproductive phenology emerges as a byproduct of selection for efficient tracking of spatio-temporal resource peaks, with the mating area corresponding to the region exhibiting the highest annual resource peak. Under the site-dependent hypothesis, reproduction is constrained by the spatial location of suitable breeding grounds. In this case, individuals aggregate in the mating area even if it is not the most productive location overall, but the mating season is expected to match the local peak in productivity within the mating area, resulting in synchronization between mating timing and the resource peak at the occupied site while other areas may remain more productive. Under the mixed hypothesis, individuals track peaks in resource availability and occupy high-productivity areas. However, annual peaks are comparable across sites. This suggests that spatial resource tracking and site dependence act as concurrent primary drivers of migration.

However, migration is also often regarded as an adaptive resource-tracking strategy in seasonal environments, with individuals that move to exploit spatio-temporal peaks in resource availability (Van der Graaf et al., 2006; Dingle et al., 2007; Somveille et al., 2018). This is, for example, and preeminently, the case for large herbivore mammals inhabiting seasonal environments, with breeding strategies (long gestation, precocial calves) that make the availability of trophic resources the only truly limiting factor (Holdo et al., 2009; Merkle et al., 2016; Purdon et al., 2018). Under this trophic resource-driven mechanism, reproductive phenology is expected to evolve purely from selection favoring effective resource tracking. In this framework, migratory behavior is indeed expected to emerge primarily from the need to track shifting resource peaks across space and time, with reproductive timing adjusting accordingly such that breeding occurs in the area experiencing the highest annual trophic resource peak among those visited throughout the migratory cycle (Figure 1).

A third scenario may emerge when these alternative mechanisms become difficult to disentangle. In such cases, migration may simultaneously reflect the need to track seasonal resource dynamics and the constraint imposed by the access to suitable breeding sites, with neither process acting as a clearly dominant driver. Under this mixed mechanism, individuals are still expected to exploit spatially and temporally variable resource peaks and preferentially occupy highly productive areas throughout the annual cycle. However, the breeding area does not necessarily correspond to the site exhibiting the highest annual trophic resource peak among those visited while migrating. Instead, comparable resource peaks across sites may allow breeding site access and resource tracking to jointly shape migratory behavior and reproductive phenology, suggesting that site dependence and trophic optimization operate as concurrent primary drivers of migration (Figure 1).

Although different mechanisms may prevail across ecological contexts, empirical tests simultaneously evaluating the spatial structure of breeding sites and the seasonal dynamics of trophic resources remain almost completely restricted to birds and large mammals (Soriano-Redondo et al., 2020). Here, we take advantage of a unique natural system to evaluate competing hypotheses regarding these mechanisms in a reptile species exhibiting partial migration. The Galápagos pink land iguana, *Conolophus marthae* (Gentile & Snell, 2009; hereafter also referred to as “pink iguana”), is a critically endangered species (Gentile, 2012; Gentile et al., 2016) of about 200 adult individuals (Garizio et al., 2024) endemic to the north-western slope of Wolf Volcano, Isabela Island, Galápagos (Gargano et al., 2026). Previous research and long-term field observations suggest that pink iguanas congregate near the rim of Wolf Volcano (*ca.* 1700 m a.s.l.) between March and July for courtship and mating (Colosimo et al., 2022; Onorati et al., 2016). After mating, females move up to 2 km inside the caldera to nest (Gargano et al., 2024). At the end of the reproductive season, a large proportion of individuals disperse downslope (Appendix S1). Interestingly, a similar pattern of seasonal aggregation and dispersal has been described for the congeneric Galápagos land iguana (*C. subcristatus;* hereafter also referred to as “yellow iguana”) on Fernandina Island (Werner, 1982). *Conolophus subcristatus* also occurs in syntopy with *C. marthae* on Wolf Volcano, and field observations indicate that yellow iguanas may adopt the same strategy on Wolf Volcano as well (Kumar et al., 2020). Such a shared migratory behavior may either represent a plastic response to similar local conditions, a genetically-determined ancestral trait or a strategy evolved independently under similar environmental constraints, making this system particularly suitable for evaluating the main drivers of partial migration.

Although in *C. marthae* such a seasonal pattern has been previously documented in a single individual (Colosimo et al., 2022), a formal, multi-individual assessment is still lacking, and the ecological and evolutionary bases of this migration remain completely unexplored. Here, we provide the first comprehensive analysis of the species’ movement patterns based on multiple tracked individuals. Using this system, we further test whether the observed movements are primarily driven by the spatial constraint imposed by the breeding sites or reflect a strategy to track predictable seasonal changes in environmental conditions. Because *C. marthae* is an ectothermic species, we also evaluate an alternative explanation whereby seasonal movements may reflect the need to track suitable thermal conditions across the altitudinal gradient, with migratory individuals potentially gaining fitness advantages by shifting to lower elevations during the dispersal period to optimize physiological performance and enhance survival, as frequently happens in reptiles (Harvey & Larsen, 2020).

## Materials and Methods

### Data collection

Between 2019 and 2022, custom GPS-Wireless Sensor Nodes (WSNs; Loreti et al., 2019; Loreti et al., 2020) were attached to a total of 42 adult pink land iguanas (17 males and 25 females; Appendix S2) captured across the entire species altitudinal range (approx. 600– 1600 m a.s.l.; Figure 2). Research activities, including animal capture and the deployment of WSNs, were authorized by the Galápagos National Park Directorate (GNPD) through Scientific Research Permits No. PC-92-19, PC-04-21, and PC-50-22. All procedures were conducted in accordance with the regulations governing scientific research within the Galápagos Protected Areas. While subsets of these movement data contributed to previous technological validations and ecological studies (Loreti et al., 2020; Colosimo et al., 2022; Gargano et al., 2024), the present work integrates the full available dataset to provide a comprehensive, population-level assessment of seasonal migration and its environmental drivers.

**Figure 2.**
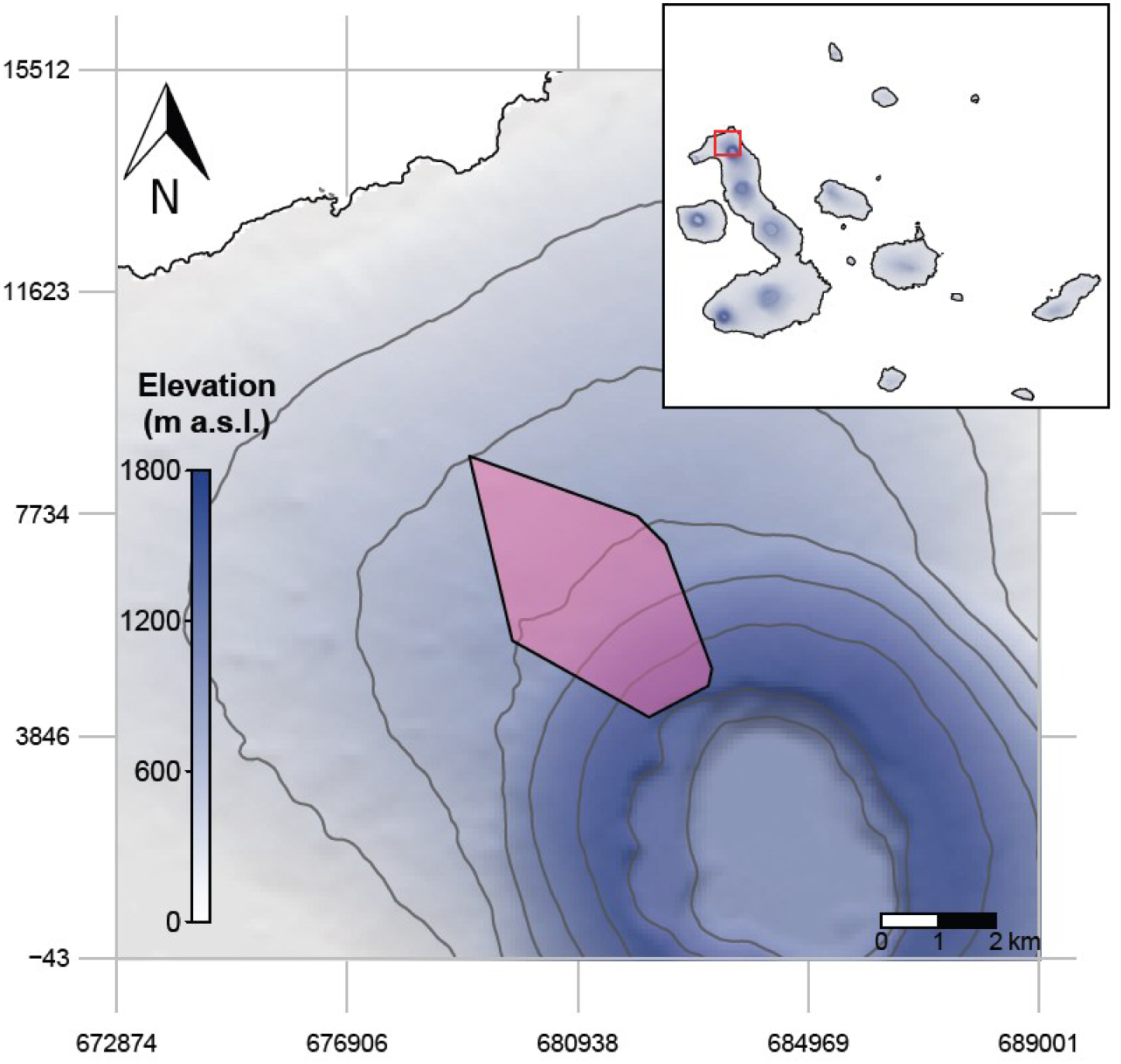
Minimum Convex Polygon (MCP) based on the capture locations of the 42 individuals of *C. marthae* tracked on Wolf Volcano between 2019 and 2023. Individual capture locations are not shown to avoid disclosing the exact locations of this critically endangered species. Contour lines represent 300 m interval change in elevation. Map projection: UTM Zone 15N, WGS84.

### Microclimate and productivity reconstruction

Within an established conservation framework for *C. marthae* (Rueda et al., 2023), the deployment of 10 meteorological stations (HOBO® Micro Station H21-USB) across the entire altitudinal range occupied by *C. marthae* was completed by 3 December 2022 (Appendix S3; Appendix S4). We leveraged these data to generate spatio-temporally continuous predictions of mean air temperature across the entire study area and throughout the full annual cycle using Generalized Additive Models (GAMs). As a result, we produced a complete series of daily raster layers at 250 m resolution. In parallel, the same methodological pipeline was applied to produce spatio-temporal predictions for mean relative humidity, total precipitation, and mean ultraviolet radiation. Seasonal patterns of vegetation productivity were characterized using 16-day composite layers of the Normalized Difference Vegetation Index (NDVI) for the period 2000–2024, derived from the MODIS MOD13Q1 product (Collection 6.1; Didan, 2021). A harmonic regression approach based on truncated Fourier series (Menenti et al., 1993; Moody & Johnson, 2001; Beck et al., 2006) was applied to reconstruct daily NDVI time series for each 250 m pixel within the study area. Detailed procedures regarding data filtering, model fitting, validation and prediction for all environmental variables are provided in the Supplementary Materials (Appendix S5; Appendix S6; Appendix S7).

Both air temperature and NDVI predictions were interpreted as representative of a typical year for the restricted area on Wolf Volcano. Galápagos climate is indeed characterized by a regular seasonal cycle with limited inter-annual variability, except for major El Niño– Southern Oscillation (ENSO) events (Nieuwolt, 1991; Trueman & D’Ozouville, 2010). Accordingly, modelling climate using one full seasonal cycle and deriving NDVI trajectories from multi-year composites should capture the dominant and recurring environmental conditions characterizing the study area.

### Movement reconstruction and migration timing estimation

Location data from the 42 individuals tracked between 2019 and 2023 were subjected to a filtering procedure to ensure spatial and temporal reliability. First, we kept only positions displaying suitable accuracy values by checking the Horizontal Dilution of Precision (HDOP), and values > 1.4 were excluded (Loreti et al., 2020). To avoid over-representation of repeatedly sampled locations, a single location per day for each individual was considered, and the remaining data were then thinned by retaining a single random location per individual within each 5-day temporal window and cell. Finally, individuals with fewer than 15 retained locations after filtering were excluded from subsequent analyses.

To quantify iguanas’ altitudinal movement patterns, we adopted the nonlinear model-based approach proposed by Bunnefeld et al. (2011) but adapted to directly model elevation rather than net squared displacement as a function of time. This allowed us to capture directional movements along the elevation gradient, explicitly representing altitudinal displacement processes. For each individual, elevation trajectories were fitted using a set of models representing alternative movement behaviors, including stationary behavior (constant model), monotonic altitudinal shifts (single sigmoid), and bidirectional movements (double sigmoid). Models were fitted to individual trajectories using nonlinear least squares and compared using the Akaike’s Information Criterion (AIC). The model with the lowest AIC value was selected as the best-supported movement model for each individual. For pink iguanas displaying altitudinal displacement, onset and termination of movement phases were derived from the model coefficients. Specifically, the start of displacement was defined as the earliest point at which the rate of altitudinal change exceeded 10% of the maximum observed rate, while the end of displacement corresponded to the earliest point at which the rate of change declined below the same threshold.

To characterize population-level seasonal movement, we aggregated individual displacement timing metrics derived from the best-supported models. Population patterns were summarized by calculating the median day of year (DOY) of each transition across individuals. When sample size allowed, transition dates were summarized separately for males and females to account for potential sex-specific differences in movement phenology. These sex-specific medians were used to reconstruct the general annual cycle of the migratory population, specifically defining a mating season, a nesting season, and a dispersal season.

### Defining functional areas, aggregation patterns, and associated environmental variables

To characterize the environmental conditions associated with the seasonal movement of *C. marthae*, we first identified two functional areas: a mating area and a dispersal area. These areas were defined using 427 capture locations collected during long-term monitoring of the species between 2005 and 2022. The spatial extent of each area was estimated by computing the 95% kernel utilization distribution (KUD, Calenge, 2024) on subsets of these occurrence records, filtered according to the estimated population-level seasonal movement (see above) and altitudinal thresholds. Specifically, the mating area was defined using capture records (*n* = 107) occurring during the mating season and located above 1500 m a.s.l. Based on these records, we estimated the spatial extent of this area to be 2.05 km^2^. The dispersal area was defined using capture records (n = 36) collected during the dispersal season and occurring below 1500 m a.s.l., resulting in a total area of 25.88 km^2^. For each functional area in each season differentially defined by sex, we calculated the daily mean NDVI and temperature values to obtain a time series of environmental conditions associated with each area and each season.

At the same time, to quantify the spatial aggregation pattern associated with the two seasonal phases, we compared the spatial distribution of capture locations between the mating and dispersal seasons. Because our aim was to assess aggregation at the scale of the population home range rather than within season-specific functional areas, we defined a common spatial reference area by estimating the 95% kernel utilization distribution (KUD)using all available capture locations (Calenge, 2024). Capture locations were then assigned to the mating and dispersal periods according to the seasonal classification described above. To obtain comparable estimates between periods, the mating-period dataset was randomly subsampled to match the number of capture locations available for the dispersal season. For each season, we quantified the spatial aggregation of individuals using two complementary descriptive metrics. First, we calculated the 100% minimum convex polygon (MCP; Calenge, 2024) area as a measure of the spatial extent occupied by individuals during each period. Second, we calculated the nearest neighbor index (NNI) to quantify local spatial aggregation relative to the expected distribution under complete spatial randomness. NNI was computed as the ratio between the observed mean nearest-neighbor distance and the expected mean nearest-neighbor distance given the number of locations and the area of the common spatial window (Clark & Evans, 1954). Values of NNI < 1 indicate increasing levels of spatial aggregation, whereas values approaching 1 indicate a random spatial distribution. Together, MCP and NNI provided complementary measures of spatial aggregation, describing respectively the extent of occupied space and the degree of local clustering of capture locations.

### Testing alternative hypotheses about migratory strategy evolution

To test alternative hypotheses explaining the evolution of seasonal migration in *C. marthae*, we analyzed how environmental conditions differed between functional areas between the mating and the dispersal season, by fitting separate linear models with NDVI and temperature as the respective response variables and area identity, season, sex and their interactions as predictors.

### Computational environment and data analysis

All data processing and statistical analyses were conducted in R v4.2.2 (R Core Team, 2022) using RStudio v2024.9.1.394 (R Studio Team, 2024), while the extraction and preprocessing of NDVI data were performed in Google Earth Engine (GEE; https://earthengine.google.com/). Details of the packages used, and the analytical procedures implemented are provided in the corresponding analysis scripts, which will be made publicly available through the Data Availability Statement to ensure transparency and reproducibility.

## Results

### Migratory patterns and timing

After data filtering, we reconstructed altitudinal trajectories and estimated migration timing for 22 of the 42 tracked individuals (7 males and 15 females; Figure 3). Model selection revealed heterogeneous movement strategies within the monitored population: altitudinal paths of 9 individuals were best described by a double sigmoid model, indicating two distinct phases of altitudinal displacement; 7 by a single sigmoid model, reflecting a single seasonal shift; and 6 by a constant model, with no detectable directional altitudinal movement during the study period (Figure 3; Appendix S8). Altitudinal displacement was therefore detected in 16 out of 22 individuals. Movements toward the mating area were concentrated between December and February (DOY 340-46) for males (*n* = 4) and January and April (DOY 2-97) for females (*n* = 4) (Figure 3). Movements to the nesting area were detected for 10 females, from April and May (DOY 108-144). For 6 of these females, the departure from the nesting area and the return to the rim was also recorded, and occurred between May and June (DOY 124-169). For only one male individual (WSN_48, Appendix S9), continuous tracking was available from July through November. This animal remained on the volcano rim throughout the whole period. For additional four individuals (two females: WSN_30 and WSN_43 and two males: WSN_36 and WSN_41), position data were available from September through November. Among these individuals, WSN_36, a male, descended to the dispersal area in October (DOY 278-286), while the females and the remaining male stayed on the volcano rim throughout the whole period (Figure 3; Appendix S9; Appendix S10). Population-level migratory timing was summarized using the median DOY of individual transition dates (Figure 3; Figure 4a). Males exhibited two major seasonal movements, consisting of arrival at the mating area at the end of December (DOY 353) and descent to the dispersal area in October (DOY 286) of the following year (although this timing was estimated based on a single individual). In females, the annual cycle included arrival at the mating area at the beginning of February (DOY 36), 48 days later than males, followed by movements to and from the nesting area in May (DOY 125-146). Descending movements to dispersal areas were not detected in the tracking dataset for females. However, females are regularly captured at lower elevations during the dispersal season (Appendix S1), implying that at least some of them also experience a downward migration at some time between July and November. In the following, we tentatively assumed that the start of the dispersal season for females is the same as that estimated for males.

**Figure 3.**
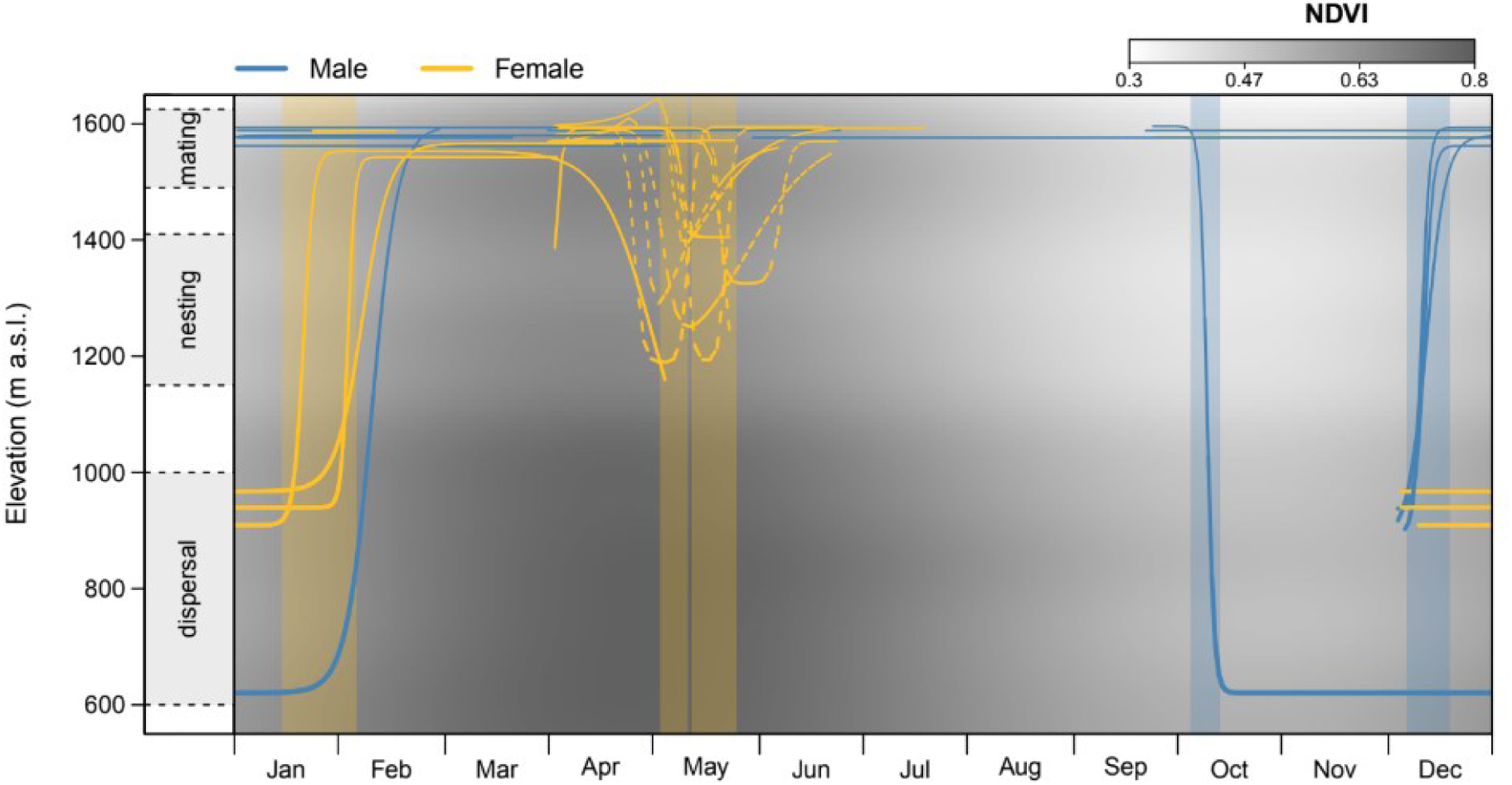
Fitted altitudinal movement patterns of the 22 tracked individuals of *C. marthae* retained after data filtering. Lines thickness is inversely scaled to elevation, and line color denotes sex. Dashed lines highlight the nesting migration for females. Vertical semi-transparent rectangles indicate the estimated population-level migration timing events for both sexes. The background color shows the seasonal variation of mean NDVI across the altitudinal gradient occupied by the tracked individuals. Reference bars on the left indicate the approximate altitudinal ranges of mating, nesting, and dispersal areas.

**Figure 4.**
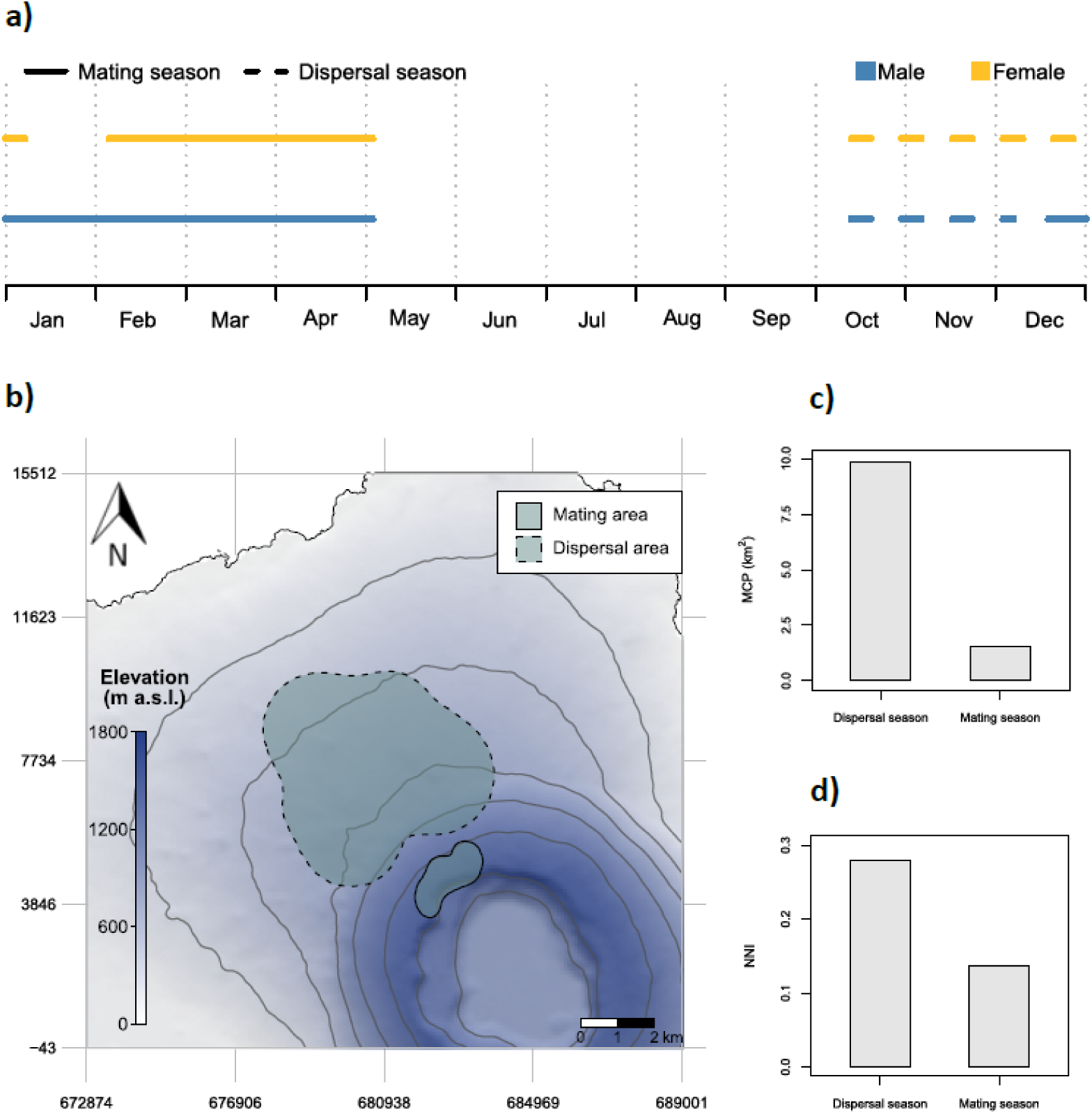
Main features of the partial migration of *Conolophus marthae*. Panels show *a)* the annual timeline of migratory phases for males and females, *b)* the spatial distribution of the functional areas used by the species across seasons, *c)* the 100% Minimum Convex Polygon (MCP, km^2^) area occupied by capture locations during the mating and dispersal seasons, as a descriptive measure of the spatial extent occupied by individuals, and *d)* the Nearest Neighbor Index (NNI) for the two seasonal phases, describing the degree of local spatial aggregation of capture locations relative to a random spatial distribution. Lower NNI values indicate stronger spatial aggregation.

### Seasonal differences in spatial aggregation

Capture locations showed stronger spatial aggregation during the mating season. The 100% MCP estimated an occupied area of 1.563 km² during the mating season and 9.847 km² during the dispersal season (Figure 4c; Appendix S11), indicating a substantially more restricted spatial concentration of individuals during mating. Similarly, the NNI showed strong clustering in both periods (Figure 4d; Appendix S11), with lower values during mating (NNI = 0.136) compared to dispersal (NNI = 0.280).

### Drivers of migration

The linear model with NDVI as the response variable revealed significant differences in productivity between functional areas (Appendix S12). The dispersal area was generally more productive than the mating area throughout the year (Figure 5a). However, the mating season coincided with the local peak of vegetation productivity within the mating area. In particular, females reached the mating area when productivity was close to its seasonal maximum, while males arrived earlier, occupying the mating area when productivity had not yet reached the local peak (Figure 4a; Figure 5a).

**Figure 5.**
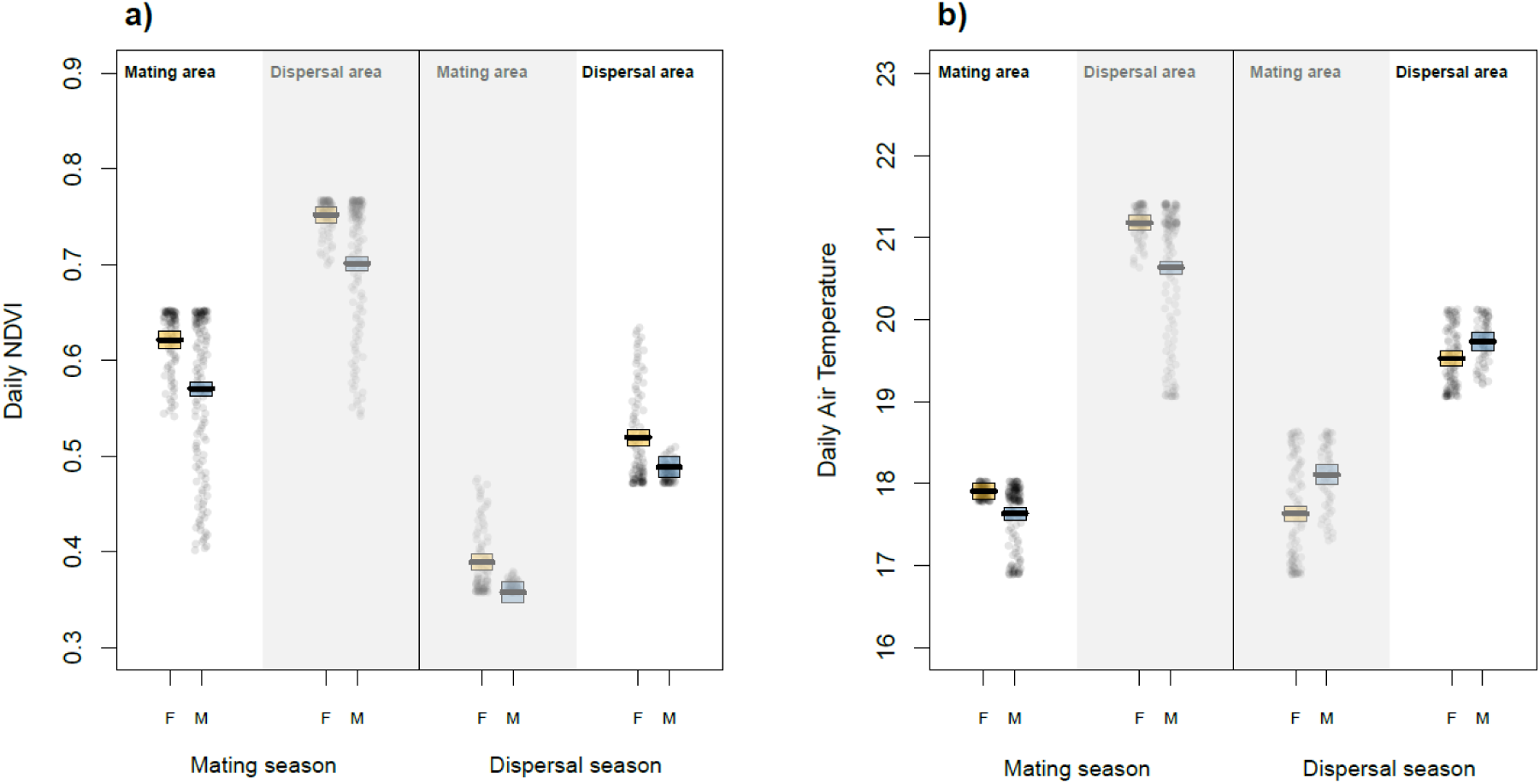
Daily values of the Normalized Difference Vegetation Index (NDVI) and air temperature measured in the mating and dispersal areas of *C. marthae*. Values are shown separately for the mating season and the dispersal season and are further grouped by sex (F = females, M = males). Each point represents a daily mean NDVI (a) or air temperature (b) for the corresponding area and period. Note that the start and end dates of the mating and dispersal seasons differ between males and females, while functional areas are defined at the population level. The different values for males and females, therefore, depend on the differences between the time intervals spent by each sex in each area (Figure 4a). For each season, the functional area actually occupied by pink iguanas (e.g., the mating area during the mating season) is shown with standard colours, whereas the alternative functional area (i.e., the area not occupied during that season) is displayed with faded colours on a grey background, highlighting where individuals were versus where they could potentially have been. Boxes superimposed on the raw data summarize the fitted model predictions: the central horizontal line represents the model estimate, and the upper and lower box edges indicate the 95% confidence interval.

Air temperature exhibited the expected altitudinal gradient, with warmer conditions in the low-elevation dispersal area and cooler temperature near the rim of the volcano (Figure 5b), although the magnitude of these differences remained relatively small (*ca.* 3°C in the mating season and *ca.* 2°C in the dispersal season). Seasonal variation in temperature was also detected but was generally modest, being essentially null in the mating area and < 2°C in the dispersal area (Figure 5b; Appendix S12).

## Discussion

This study provides a formal quantitative assessment of the relative roles of spatial constraints associated with critical life-history sites and trophic resource availability in shaping partial migration in a large terrestrial reptile. Using the Galápagos pink land iguana as a natural model system, our results also represent the first population-level evaluation of seasonal movements for this species. Integrating tracking data from 22 adult individuals with reconstructed high-resolution environmental variables, we provide a key empirical contribution toward understanding the evolutionary factors shaping migration in this taxon. Our results reveal a clear pattern of partial migration along the altitudinal gradient of Wolf Volcano and provide empirical support for the hypothesis that migration in this species is primarily driven by access to spatially restricted breeding sites rather than by constraints related to spatio-temporal variation in resource availability.

A large proportion of individuals exhibits directional movements toward a restricted high-elevation mating area (Figure 3), with males arriving in late December and females arriving around two months later (Figure 4a). This upward migration involves substantial elevational displacements, ranging between 600 to 1000 m (Figure 3; Appendix S9; Appendix S10). Although we do not have evidence of iguanas remaining in the dispersal area during the mating season, we cannot exclude that some individuals, potentially young subadults, may remain at lower elevation throughout the year. In females, the ascent phase is followed by a distinct nesting migration occurring between April and June, consisting of movements from the mating area toward nesting grounds within the Wolf volcano caldera and subsequent returns to the rim (Figure 3; Figure 4a). This nesting migration has been previously described in a subset of individuals (Gargano et al., 2024) and now confirmed and extended by the analysis of additional females (Appendix S9; Appendix S10). After the reproductive period, individuals move toward lower elevations, marking the onset of the dispersal phase (Figure 3). This pattern is supported by yet unpublished mark-recapture data showing that individuals captured at lower elevations during the dispersal season are later recaptured near the rim during the mating season, although descending movements are documented by tracking data for only one individual in the dataset (Figure 3). Notably, at least one individual (WSN_48, Appendix S9) remained at high elevation throughout the entire annual cycle, confirming that this system represents a case of partial migration, in which altitudinal shifts are performed by only a subset of individuals. At the population level, these movements define a cyclical pattern of seasonal aggregation at high elevation during reproduction and broader spatial redistribution across lower elevations during the non-reproductive season (Figure 4b-c).

Our analysis shows that the mating area does not coincide with the most productive location available within the species range (Figure 5a; Appendix S12). Indeed, the low-elevation dispersal area remains more productive throughout the year (Figure 5a; Appendix S12), indicating that individuals do not simply track resource peaks in space and time. Instead, migration appears to be primarily driven by the spatial constraint imposed by the location of suitable breeding grounds. However, the mating season coincides with the local peak in vegetation productivity of the mating area (Figure 5a), suggesting that the timing of reproduction has evolved to align with optimal local conditions, consistent with the idea that resource dynamics act as a secondary selective pressure shaping reproductive phenology rather than driving migration per se. This pattern closely matches the conceptual expectations of the site-dependent hypothesis and contrasts with predictions of the resource-driven hypothesis (Figure 1), under which individuals would be expected to occupy the most productive area available at any given time (Dingle et al., 2007; Somveille et al., 2018).

More broadly, our findings align with theoretical models proposing that partial migration can evolve when access to breeding sites represents the ultimate limiting resource (Lundberg, 1987; Kokko & Lundberg, 2001). In such systems, individuals may accept suboptimal foraging conditions to secure reproductive opportunities, resulting in a spatial decoupling between optimal feeding and breeding habitats (Lundblad & Conway, 2020). The pronounced sex-specific differences in migratory timing observed in our study further support this interpretation, providing additional insight into the drivers of this behavioral strategy. Males indeed reached the mating area earlier than females, arriving when local productivity peak had not yet been reached (Figure 4a; Figure 5a; Appendix S12). This anticipatory movement indicates that males prioritize early access to mating sites, likely to establish and defend territories before the onset of the mating season (Werner, 1982). In this view, males would accept the cost of leaving more productive lowland areas at a time when resource availability is increasing (Figure 3; Figure 5), highlighting a trade-off between foraging opportunities and reproductive success (Lundberg, 1987). In contrast, females arrived at the mating area when vegetation productivity was close to its seasonal maximum, reasonably to optimize energy allocation prior to reproduction and synchronize their arrival with favorable local conditions, since food intake in this phase may strongly affect their reproductive output (Bonnet et al., 2001). These sex-specific strategies are consistent with movement patterns described in *C. subcristatus* by Werner (1982) on Fernandina Island, where individuals aggregate in restricted mating areas coming from around 3 km away, with males that arrive earlier to establish their territories. After mating, males occupying areas with insufficient local productivity disperse, whereas females migrate for more than 15 km to reach the rim of La Cumbre volcano and nest inside the caldera. In *C. marthae*, both sexes perform migrations of up to 5 km to reach the mating area (Colosimo et al., 2022), while females later undertake shorter nesting migrations of about 1 km (Gargano et al., 2024). Despite these differences in magnitude, the overall structure of the system remains analogous. Our results thus highlight the potential of the Galápagos land iguanas clade as a model for the evolution of migratory behavior in reptiles.

Interestingly, temperature does not appear to play a major role in shaping migration in this system (Figure 5b). Despite the wide altitudinal range covered by pink iguanas in their migratory movements (*ca.* 600 - 1600 m asl), temperature differences between functional areas were relatively small (*ca.* 2°C) and did not show consistent seasonal or sex-specific patterns that might explain migration timing (Figure 5b). This suggests that seasonal movements are unlikely to be driven by direct physiological constraints related to thermoregulation. This result contrasts with a recurrent interpretation of patterns observed in other reptiles, which sees altitudinal migration as a strategy to maintain suitable body temperatures (Harvey & Larsen, 2020) and highlights that different ecological contexts can favor distinct movement strategies.

Our results also have direct conservation implications for the critically endangered *C. marthae*. Recent species distribution modelling has identified potential receptor sites for the establishment of a second population, providing an important framework for future *ex-situ* conservation actions (Gargano et al., 2026). However, our results highlight the importance of further evaluating candidate areas in relation to the specific reproductive resources and spatial requirements that underpin the species migratory cycle. Since seasonal movements are mainly driven by access to restricted breeding areas, successful translocation will require the presence of suitable nesting and mating grounds to support reproductive aggregation and recruitment. Moreover, all the identified candidate areas overlap with the distribution of *C. subcristatus* (Gargano et al., 2026), suggesting that competitive interactions should be explicitly considered when assessing translocation feasibility. Both the extensive overlap in the two species’ isotopic niches (Gargano et al., 2022) and their demographic histories (Paradiso et al., 2025) strongly suggest that competitive exclusion may have limited the distribution of *C. marthae*. Our results suggest that competitive dynamics could be further exacerbated during the reproductive season. Since pink iguanas occur at high density in the mating area during its productivity peak, overlaps in habitat use with *C. subcristatus* at this site may lead to intensified competition for the limited trophic resources available during congregation. Moreover, while nesting sites on Wolf Volcano appear to be distinct (Kumar et al., 2020; Gargano et al., 2024), the potential for competition for suitable nesting grounds in new receptor areas remains an open question, representing a significant risk for translocation success. In addition, the observed link between reproduction and vegetation greenness highlights the species’ vulnerability to environmental shifts. The projected alterations in Galápagos productivity caused by climate change (Charney et al., 2021) suggest that variations in timing and magnitude of vegetation green-up may represent a serious threat by disrupting the synchronization of the species’ migratory phenology. Given that pink iguanas congregate in the mating area during the local productivity peak (Figure 4; Figure 5a), any phenological mismatch could jeopardize the already limited reproductive success (Garizio et al., 2024).

Finally, our study provides more general resources for conservation in the Galápagos: we expect that, beyond the pink iguana, the high-resolution (250m) environmental rasters we produced will prove a valuable resource for the management and protection of the Wolf Volcano ecosystem. By capturing fine-scale microclimates missed by global models, they facilitate the study of sympatric species like giant tortoises, enabling the identification of key habitats and seasonal vulnerabilities essential for effective conservation planning.

## Acknowledgements

We thank the Galápagos National Park Directorate (GNPD) for supporting this research within the framework of the long-term collaboration with the University of Rome “Tor Vergata” aimed at the conservation of Galápagos land iguanas. We are grateful to the park rangers for their essential assistance during field activities. Fieldwork and WSN development was supported by Friends of Galápagos Foundation, the San Diego Zoo Wildlife Alliance [research grant to GC and GG], the International Iguana Foundation [grant 2018-2019 to GG], the Mohamed bin Zayed Species Conservation Fund [grant number 12254183 and 182518617 to GG], the National Geographic Fund [grant number NGS-68399C-20 to GG], the Foundation Segré Conservation Action Fund [grant 2022-2023 to GG] and the “Uncovering Excellence” grant from the University of Rome “Tor Vergata”. Additional support for the 2021 expedition was provided by Canal+ Docs. Research activities were conducted under permits issued by the Galápagos National Park Directorate (PC-92-19, PC-04-21, and PC-50-22).

## Author Contributions

**Marco Gargano:** Conceptualization, Formal analysis, Investigation, Data Curation, Writing - Original Draft. **Lorenzo Garizio:** Conceptualization, Formal analysis, Investigation, Data Curation, Writing - Original Draft. **Giuliano Colosimo:** Conceptualization, Investigation, Data Curation, Writing - Review & Editing. **Pierpaolo Loreti:** Resources, Writing - Review & Editing; **Alexandro Catini:** Resources, Writing - Review & Editing. **Lorenzo Bracciale:** Resources, Writing - Review & Editing. **Massimiliano De Luca:** Resources, Writing - Review & Editing. **Gregory A. Lewbart:** Investigation, Writing - Review & Editing. **Christian Sevilla:** Resources, Investigation. **Glenn P. Gerber:** Funding acquisition, Writing - Review & Editing. **Paolo Gratton:** Conceptualization, Supervision, Writing - Review & Editing. **Gabriele Gentile:** Conceptualization, Investigation, Data Curation, Resources, Writing - Review & Editing, Supervision, Project administration, Funding acquisition.

## Data availability statement

The data supporting the findings of this study will be archived in Figshare and made available upon acceptance of the manuscript. To protect the critically endangered *Conolophus marthae*, original GPS coordinates and fine-scale locations of individuals will not be publicly released. The repository will instead provide processed datasets that preserve the structure of the analyses while preventing the disclosure of sensitive localities.

