## Supplementary material for "Why they move: breeding-site availability drives partial migration in a large terrestrial reptile"

SUPPLEMENTARY MATERIALS

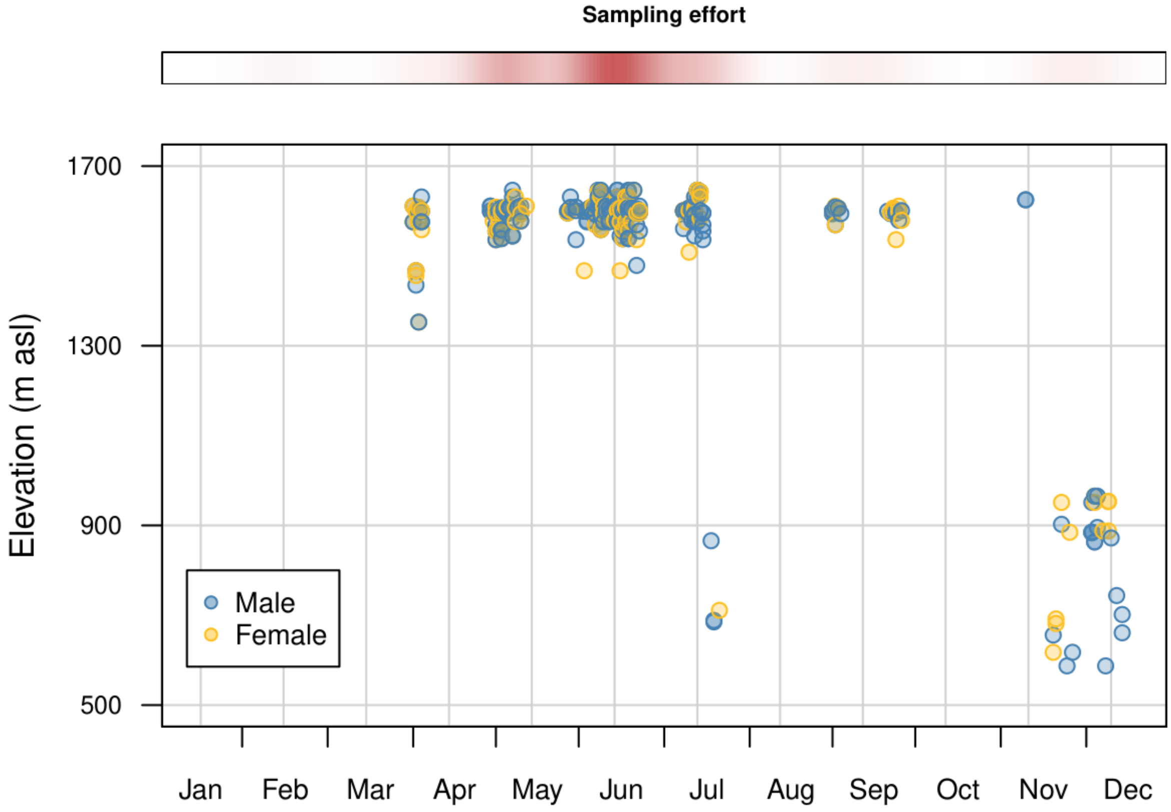


**Appendix S1. Altitudinal and seasonal distribution of *C. marthae* captures based on long-term monitoring data (2005–2025).** Points represent capture events for males (in blue) and females (in orange). The upper heatmap represents the density of sampling effort throughout the year.

**Appendix S2. Summary of deployment information and record history of the GPS-Wireless Sensor Nodes (WSNs) attached to pink iguanas between 2019 and 2022.** For each individual, the table reports WSN ID, sex of the individual, capture date, date of last recorded location, total number of raw records collected, and number of filtered records retained after data cleaning procedures. Filtered records represent the subset of locations used in the movement analyses after applying quality and subsampling criteria (see Methods for details).

| WSN ID | Sex | Capture | Last record | Raw records (n) | Filtered records (n) |
|---|---|---|---|---|---|
| **WSN_13** | F | 25/09/2019 | 18/10/2019 | 65 | / |
| **WSN_16** | F | 04/04/2021 | 14/06/2021 | 50 | / |
| **WSN_18** | F | 02/04/2021 | 24/06/2021 | 268 | 26 |
| **WSN_20** | M | 23/09/2019 | 23/09/2019 | 1 | / |
| **WSN_23** | F | 04/04/2021 | 16/05/2021 | 111 | 17 |
| **WSN_24** | M | 21/09/2019 | 06/10/2019 | 81 | / |
| **WSN_28** | M | 02/04/2021 | 25/06/2021 | 184 | 28 |
| **WSN_29** | F | 03/04/2021 | 24/05/2021 | 94 | 22 |
| **WSN_30** | M | 26/09/2019 | 04/05/2020 | 656 | / |
| **WSN_31** | F | 05/04/2021 | 07/06/2021 | 231 | 30 |
| **WSN_32** | M | 24/09/2019 | 29/09/2019 | 43 | / |
| **WSN_33** | F | 04/04/2021 | 18/08/2021 | 179 | 20 |
| **WSN_36** | M | 24/09/2019 | 02/03/2020 | 355 | 45 |
| **WSN_38** | F | 26/09/2019 | 16/02/2020 | 321 | 34 |
| **WSN_39** | F | 03/04/2021 | 06/03/2023 | 467 | 38 |
| **WSN_40** | F | 04/04/2021 | 26/06/2021 | 256 | 30 |
| **WSN_41** | M | 22/09/2019 | 28/01/2020 | 140 | 17 |
| **WSN_43** | F | 25/09/2019 | 27/12/2019 | 252 | 28 |
| **WSN_46** | F | 02/04/2021 | 22/06/2021 | 87 | 18 |
| **WSN_47** | F | 03/04/2021 | 25/05/2021 | 71 | / |
| **WSN_48** | M | 02/04/2021 | 22/03/2022 | 744 | 74 |
| **WSN_50** | M | 23/09/2019 | 12/10/2019 | 86 | / |
| **WSN_51** | M | 04/12/2022 | 05/05/2023 | 534 | 40 |
| **WSN_52** | F | 23/09/2019 | 23/09/2019 | 7 | / |
| **WSN_53** | F | 02/04/2021 | 23/06/2021 | 127 | 26 |
| **WSN_54** | F | 03/04/2021 | 22/03/2022 | 205 | 19 |
| **WSN_55** | F | 22/09/2019 | 22/09/2019 | 1 | / |
| **WSN_58** | F | 02/04/2021 | 10/04/2021 | 17 | / |
| **WSN_60** | F | 22/09/2019 | 26/09/2019 | 30 | / |
| **WSN_61** | F | 24/09/2019 | 08/10/2019 | 29 | / |
| **WSN_64** | M | 24/09/2019 | 24/09/2019 | 1 | / |
| **WSN_70** | M | 04/12/2022 | 04/05/2023 | 431 | 56 |
| **WSN_71** | M | 04/12/2022 | 04/12/2022 | 1 | / |
| **WSN_72** | F | 10/12/2022 | 07/04/2023 | 57 | / |
| **WSN_74** | F | 10/12/2022 | 05/05/2023 | 420 | 49 |
| **WSN_75** | F | 05/12/2022 | 23/04/2023 | 467 | 43 |
| **WSN_76** | M | 10/12/2022 | 10/12/2022 | 1 | / |
| **WSN_77** | M | 09/12/2022 | 05/05/2023 | 139 | / |
| **WSN_80** | M | 06/12/2022 | 04/05/2023 | 210 | / |
| **WSN_81** | M | 06/12/2022 | 04/04/2023 | 380 | 38 |
| **WSN_83** | F | 08/12/2022 | 20/04/2023 | 77 | / |
| **WSN_84** | F | 05/12/2022 | 04/04/2023 | 228 | 37 |

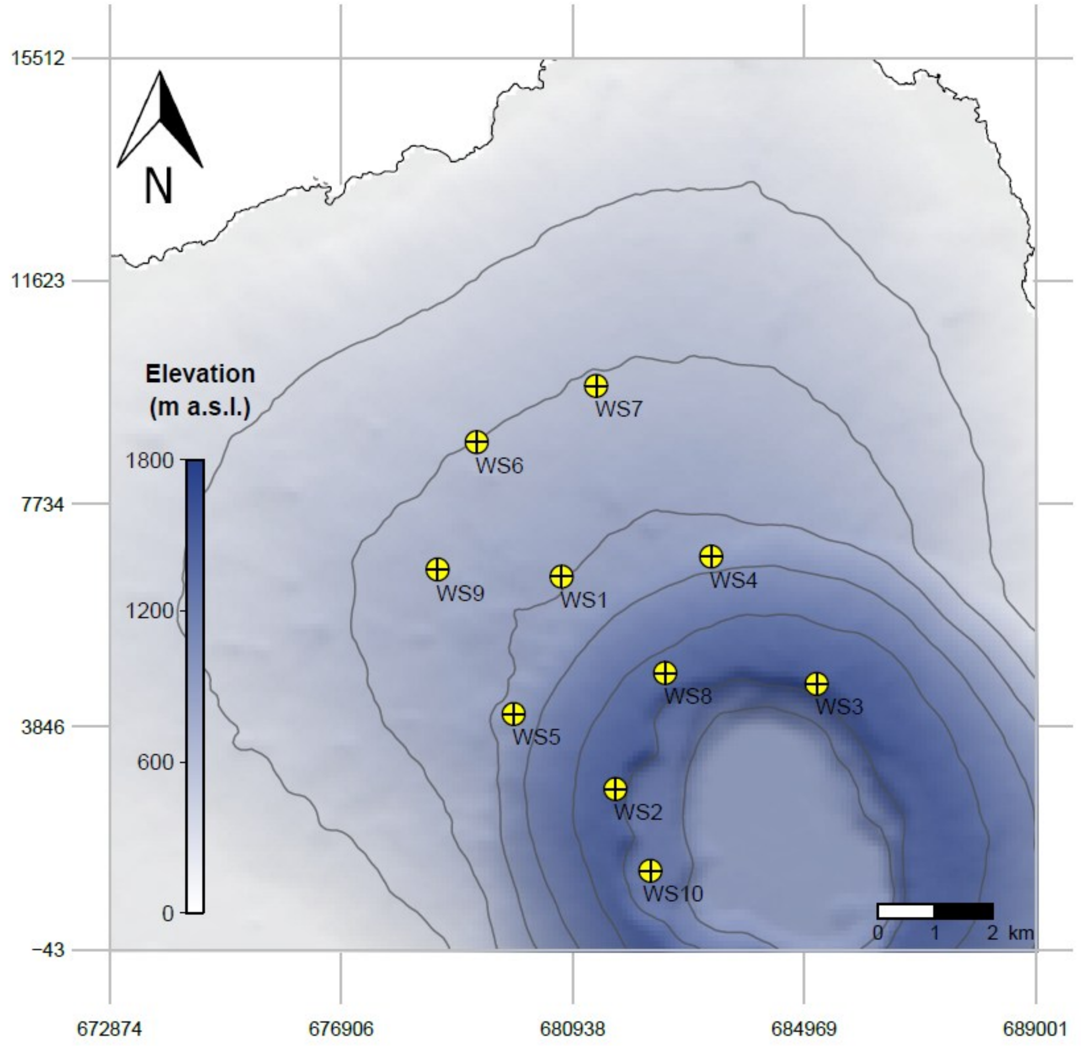


**Appendix S3. Locations of the 10 meteorological stations installed on Wolf Volcano as part of the *C. marthae* conservation framework.** Contour lines represent 300 m interval change in elevation. Map projection: UTM Zone 15N, WGS84.

**Appendix S4. Summary of records collected by the Weather Stations (WS) installed on Wolf Volcano.** For each year, the table reports the number of data collected in each month. Cells with "0" indicate that no data was collected from the station during that month, whereas cells with a slash indicate that the station had not yet been installed.

| ID | 2022 | | | | | | | | | | | |
|---|---|---|---|---|---|---|---|---|---|---|---|---|
| | Jan | Feb | Mar | Apr | May | Jun | Jul | Aug | Sep | Oct | Nov | Dec |
| WS1 | / | / | / | / | / | / | / | / | / | / | / | 2940 |
| WS10 | / | / | / | / | / | / | / | / | / | / | / | 2744 |
| WS2 | / | / | / | / | / | / | / | / | / | / | / | 2736 |
| WS3 | / | / | / | / | / | / | / | / | / | / | 133 | 2976 |
| WS4 | / | / | / | / | / | / | / | / | / | / | 153 | 2976 |
| WS5 | / | / | / | / | / | / | / | / | / | / | 39 | 2976 |
| WS6 | / | / | / | / | / | / | / | / | / | / | / | 2826 |
| WS7 | / | / | / | / | / | / | / | / | / | / | 61 | 2976 |
| WS8 | / | / | / | / | / | / | / | / | / | / | / | 2732 |
| WS9 | / | / | / | / | / | / | / | / | / | / | / | 2924 |
| | 2023 | | | | | | | | | | | |
| | Jan | Feb | Mar | Apr | May | Jun | Jul | Aug | Sep | Oct | Nov | Dec |
| WS1 | 2976 | 2688 | 2976 | 2880 | 2976 | 2880 | 2976 | 2976 | 2880 | 2976 | 2880 | 2976 |
| WS10 | 2976 | 2688 | 2976 | 2880 | 2976 | 2880 | 2976 | 2976 | 2880 | 2976 | 2880 | 2976 |
| WS2 | 2976 | 2688 | 2976 | 2880 | 2976 | 2880 | 2976 | 2976 | 2880 | 2976 | 2880 | 2976 |
| WS3 | 1510 | 0 | 0 | 0 | 0 | 0 | 0 | 0 | 0 | 0 | 0 | 0 |
| WS4 | 2976 | 2688 | 2976 | 2880 | 2976 | 2880 | 2976 | 2976 | 2880 | 2976 | 2880 | 2976 |
| WS5 | 2976 | 2688 | 2976 | 2880 | 2976 | 2880 | 2976 | 2976 | 2061 | 0 | 0 | 0 |
| WS6 | 2976 | 2688 | 2976 | 2880 | 2976 | 2880 | 2976 | 2976 | 2880 | 2976 | 1556 | 0 |
| WS7 | 2976 | 2688 | 2976 | 2880 | 2976 | 2880 | 2976 | 2976 | 2880 | 2976 | 2880 | 2976 |
| WS8 | 2976 | 2688 | 2976 | 2880 | 2976 | 2880 | 2976 | 2976 | 2880 | 2976 | 2880 | 2976 |
| WS9 | 2976 | 2688 | 2976 | 2880 | 2976 | 2880 | 2976 | 2976 | 2741 | 0 | 0 | 0 |
| | 2024 | | | | | | | | | | | |
| | Jan | Feb | Mar | Apr | May | Jun | Jul | Aug | Sep | Oct | Nov | Dec |
| WS1 | 2976 | 2784 | 2976 | 2880 | 2976 | 1904 | 0 | 0 | 0 | 0 | 0 | 0 |
| WS10 | 2976 | 2784 | 2976 | 2880 | 2976 | 2880 | 2976 | 2976 | 2880 | 2976 | 2880 | 2976 |
| WS2 | 2976 | 2784 | 20 | 0 | 0 | 0 | 0 | 0 | 0 | 0 | 0 | 0 |
| WS3 | 0 | 0 | 0 | 0 | 0 | 0 | 0 | 0 | 0 | 0 | 0 | 0 |
| WS4 | 2976 | 2784 | 2976 | 2880 | 2976 | 2880 | 2976 | 2976 | 2880 | 2976 | 2880 | 2976 |
| WS5 | 0 | 0 | 0 | 0 | 0 | 0 | 0 | 0 | 0 | 0 | 0 | 0 |
| WS6 | 0 | 0 | 0 | 0 | 0 | 0 | 0 | 0 | 0 | 0 | 0 | 0 |
| WS7 | 2976 | 2784 | 2976 | 2880 | 2976 | 2880 | 685 | 0 | 0 | 0 | 0 | 0 |
| WS8 | 2976 | 2784 | 2976 | 2880 | 2976 | 2880 | 2976 | 2976 | 2880 | 2976 | 2880 | 2976 |
| WS9 | 0 | 0 | 0 | 0 | 0 | 0 | 0 | 0 | 0 | 0 | 0 | 0 |
| | 2025 | | | | | | | | | | | |
| | Jan | Feb | Mar | Apr | May | Jun | Jul | Aug | Sep | Oct | Nov | Dec |
| WS1 | 0 | 0 | 0 | 0 | 0 | 0 | 0 | 0 | 0 | 0 | 0 | 0 |
| WS10 | 2976 | 2688 | 2976 | 2880 | 2976 | 276 | 0 | 0 | 0 | 0 | 0 | 0 |
| WS2 | 0 | 0 | 0 | 0 | 0 | 0 | 0 | 0 | 0 | 0 | 0 | 0 |
| WS3 | 0 | 0 | 0 | 0 | 0 | 0 | 0 | 0 | 0 | 0 | 0 | 0 |
| WS4 | 2976 | 2688 | 2976 | 2880 | 2976 | 137 | 0 | 0 | 0 | 0 | 0 | 0 |
| WS5 | 0 | 0 | 0 | 0 | 0 | 0 | 0 | 0 | 0 | 0 | 0 | 0 |
| WS6 | 0 | 0 | 0 | 0 | 0 | 0 | 0 | 0 | 0 | 0 | 0 | 0 |
| WS7 | 0 | 0 | 0 | 0 | 0 | 0 | 0 | 0 | 0 | 0 | 0 | 0 |
| WS8 | 2976 | 2688 | 2976 | 2880 | 592 | 0 | 0 | 0 | 0 | 0 | 0 | 0 |
| WS9 | 0 | 0 | 0 | 0 | 0 | 0 | 0 | 0 | 0 | 0 | 0 | 0 |

**Appendix S5. Microclimatic data collection and modeling.** To characterize microclimatic conditions across the study area, we installed 10 meteorological stations (HOBO® Micro Station H21-USB) along the north-western slope of Wolf Volcano between 29 November and 3 December 2022. Station deployment sites were selected to ensure representative coverage of the altitudinal range occupied by *C. marthae* (Appendix S3). Each station was equipped with sensors to measure air temperature (°C), relative humidity (%), precipitation (mm), and ultraviolet (UV) irradiance (W/m²), all manufactured by Onset Computer Corporation. Sensors were mounted at standardized heights above ground and programmed to record at 15-minute intervals, ensuring consistency across stations. The meteorological data collected by the 10 weather stations underwent a quality control procedure aimed at identifying and removing unrealistic and missing values. In detail, relative humidity values lower than 0% or higher than 100% were considered invalid and were removed. Air temperature records below −20 °C or above 50 °C were excluded, as these values exceed plausible environmental conditions for the study area (see Trueman & D'Ozouville, 2010). Negative values of precipitation were removed, as well as values exceeding 100 mm per 15-minute interval. Precipitation data from the meteorological station WS4 were excluded from further analyses due to anomalous measurements indicative of a potential sensor malfunction. WS4 systematically recorded substantially lower values than all other stations operating during the same period, with no corresponding patterns observed in neighbouring stations. Solar radiation data were filtered by removing negative values and unrealistically high radiation measurements recorded during nighttime hours (*i.e.*, values exceeding 300 W/m² between 6 pm and 6 am). Data used for predictions were restricted to the first year of data collection for each station, as several stations ceased operation after approximately one year. The filtered dataset thus included data from 4 December 2022 to 3 December 2023. From the filtered dataset, we derived daily mean air temperature and relative humidity, mean solar radiation calculated for daylight hours (6 am–6 pm), and total daily precipitation for each meteorological station.

We produced spatial predictions of microclimatic conditions for the entire study area, defined as the portion of the Wolf Volcano extending up to 1 km from the meteorological stations. The prediction area thus covered approximately 84 km² and was represented by a regular grid derived from the Shuttle Radar Topography Mission (SRTM) Digital Elevation Model (DEM; available at https://dwtkns.com/srtm30m/), which was resampled at 250 m spatial resolution. We produced climatic predictions for the study area by applying a regression-based modelling framework using R (version 4.4.2). We used Generalized Additive Models (GAMs) using the *mgcv* package (Wood & Wood, 2015) to model daily microclimatic variables as smooth functions of temporal and topographic predictors (Aalto et al., 2013). We fitted separate GAMs for daily mean air temperature, daily mean relative humidity, mean daytime solar radiation, and total daily precipitation. In all models, day of the year was included as a cyclic smooth term to capture seasonal patterns. Topographic predictors included elevation and terrain aspect, both derived from the DEM using the *raster* R package (Hijmans, 2025). The elevation was set as a linear predictor, while the aspect was included using sine and cosine transformations to account for its circular nature. For each response variable, a set of candidate models with increasing structural complexity was fitted, ranging from models including only seasonal and single topographic main effects to models incorporating interactions between seasonality and elevation and/or aspect. For daily mean air temperature and mean daytime solar radiation, we fitted GAMs with gaussian error distributions, whereas for mean relative humidity we fitted models with Beta error distribution and logit link functions and models with Tweedie error distribution and logit link functions for total daily precipitation. Model fitting was performed using restricted maximum likelihood estimation to ensure stable smoothing parameter estimation.

To assess the predictive performance of each candidate model, we applied a leave-one-out (LOO) cross-validation approach. Therefore, models were repeatedly fitted by excluding one meteorological station at a time, and predictions were generated for the excluded station. Model performance was quantified using the root mean square error (RMSE) and mean absolute error (MAE), averaged across all cross-validation folds. In addition to predictive accuracy, we explicitly evaluated spatial autocorrelation in model residuals to avoid spurious spatial structures that could bias predictions. Residuals were first averaged at the station level and then tested for spatial autocorrelation using Moran's I and Geary's C indices based on inverse-distance spatial weights. Only models showing no significant residual spatial autocorrelation ($p > 0.05$ for both indices) were considered suitable for spatial prediction. Final model selection for each microclimatic variable was based on the combination of lowest cross-validated RMSE and MAE and absence of significant spatial autocorrelation in residuals. The selected models were then used to generate daily spatial predictions across the entire study area for the considered time period.

**Appendix S6. NDVI data acquisition and predictions.** We derived spatially explicit estimates of Normalized Difference Vegetation Index (NDVI) from the Moderate Resolution Imaging Spectroradiometer (MODIS) NDVI product MOD13Q1 (Collection 6.1), which provides 16-day composite NDVI values at 250 m spatial resolution (Didan, 2021). NDVI data were accessed and processed using the Google Earth Engine (GEE) platform (https://earthengine.google.com/). The MOD13Q1 image collection was filtered to include observations acquired between 1 January 2000 and 31 December 2024 and spatially constrained to the study area extent. For each image, both the NDVI band and the associated Summary Quality Assurance (SummaryQA) layer were retained. To ensure high data reliability, only pixels flagged as "good quality" (SummaryQA = 0) were retained. NDVI values were then grouped by Julian day across all years of observation. For each pixel and each Julian date, we computed the 70th percentile of NDVI values across the full 2000–2024 time series. This percentile-based approach was adopted to characterize typical vegetation greenness while reducing the influence of residual cloud contamination and other low-quality observations. NDVI percentile layers derived in Google Earth Engine were further processed in R (version 4.4.2) using the *raster* package (Hijmans, 2025). All NDVI layers were reprojected and resampled to match the extent and resolution of the Digital Elevation Model (DEM) used in the study, using bilinear interpolation. To further reduce the influence of cloud cover on NDVI estimates, we retained only pixels supported by more than three valid observations. To reconstruct a spatially continuous daily NDVI time series representing a typical annual vegetation cycle, we applied a harmonic regression approach based on truncated Fourier series (*e.g.*, Menenti et al, 1993; Moody & Johnson, 2001). For each pixel, NDVI values were modelled as a function of time using a truncated Fourier series with a single harmonic component ($n = 1$), corresponding to an annual periodicity (period = 365 days). This choice was motivated by the marked seasonality of the Galápagos climate, which is characterized by two well-defined seasons (a warm, wetter season and a cooler dry season; Trueman & D'Ozouville, 2010). Model fitting was performed independently for each pixel using ordinary least squares. Pixels with fewer than ten valid NDVI observations across the annual cycle were excluded from modelling and assigned missing values in the predicted outputs. Using the fitted harmonic coefficients, NDVI values were predicted for each day of a standardized 365-day year, resulting in a stack of daily NDVI raster layers representing a smoothed annual trajectory of vegetation greenness. To assess the accuracy of the harmonic reconstruction, predicted NDVI values were compared against the filtered observed NDVI values for corresponding Julian day. Model performance was quantified using the mean absolute error (MAE) and root mean square error (RMSE), calculated across all pixels and Julian days with available observations. The final output consisted of 365 daily NDVI raster layers at 250 m spatial resolution, each representing the expected NDVI value for a given day of a typical year.

**References (Appendices S5 and S6)**

Aalto, J., Pirinen, P., Heikkinen, J., & Venäläinen, A. (2013). Spatial interpolation of monthly climate data for Finland: comparing the performance of kriging and generalized additive models. *Theoretical and applied climatology, 112*(1), 99-111. https://doi.org/10.1007/s00704-012-0716-9

Didan, K. (2021). MODIS/Terra Vegetation Indices 16-Day L3 Global 250m SIN Grid V061[Data set]. NASA Land Processes Distributed Active Archive Center. https://doi.org/10.5067/MODIS/MOD13Q1.061

Hijmans R (2025). raster: Geographic Data Analysis and Modeling. R package version 3.6-32, https://rspatial.org/raster

Menenti, M., Azzali, S., Verhoef, W., & Van Swol, R. (1993). Mapping agroecological zones and time lag in vegetation growth by means of Fourier analysis of time series of NDVI images. *Advances in Space Research, 13*(5), 233-237. https://doi.org/10.1016/0273-1177(93)90550-U

Moody, A., & Johnson, D. M. (2001). Land-surface phenologies from AVHRR using the discrete Fourier transform. *Remote Sensing of Environment, 75*(3), 305-323. https://doi.org/10.1016/S0034-4257(00)00175-9

Trueman, M., & D'Ozouville, N. (2010). Characterizing the Galápagos terrestrial climate in the face of global climate change. *Galápagos Research, 67*, 38–44.

Wood, S., & Wood, M. S. (2015). Package 'mgcv'. R package version, 1(29), 729.

**Appendix S7. Summary of candidate Generalized Additive Models (GAMs) and performance metrics for the spatial prediction of microclimatic variables on Wolf Volcano.** Candidate models for daily mean air temperature ($T_{mean}$), total daily precipitation ($Rain_{sum}$), mean relative humidity (RH), and mean solar radiation ($Rad_{mean}$) are presented with varying structural complexity. Predictors include day of the year (DOY), elevation (DEM), and terrain aspect (sine and cosine transformations: $asp_{sin}$, $asp_{cos}$). Predictive performance was assessed via leave-one-out (LOO) cross-validation, quantified by the Root Mean Square Error (RMSE) and Mean Absolute Error (MAE). Spatial autocorrelation in model residuals was evaluated using Moran's I and Geary's C indices; *p*-values > 0.05 indicate the absence of significant spatial autocorrelation. Final model selection was based on the best balance between minimal predictive error and non-significant residual spatial structure.

| **Temperature** | | | | |
|---|---|---|---|---|
| **Model formula** | **Moran's I (*p*)** | **Geary's C (*p*)** | **RMSE** | **MAE** |
| $T_{mean}$ ~ s(DOY, bs = "cc") + DEM + ti(DOY, by = DEM, bs = "cc") | 0.32 | 0.38 | 0.97 | 0.77 |
| $T_{mean}$ ~ s(DOY, bs = "cc") + DEM + ti(DOY, by = DEM, bs = "cc") + $asp_{sin}$ + $asp_{cos}$ + $asp_{sin}$:$asp_{cos}$ | 0.61 | 0.44 | 1.04 | 0.82 |
| $T_{mean}$ ~ s(DOY, bs = "cc") + DEM | 0.36 | 0.42 | 1.04 | 0.83 |
| $T_{mean}$ ~ s(DOY, bs = "cc") + DEM + $asp_{sin}$ + $asp_{cos}$ + $asp_{sin}$:$asp_{cos}$ | 0.61 | 0.44 | 1.12 | 0.89 |
| $T_{mean}$ ~ s(DOY, bs = "cc") + DEM + ti(DOY, by = DEM, bs = "cc") + $asp_{sin}$ + $asp_{cos}$ + $asp_{sin}$:$asp_{cos}$ + ti(DOY, by = $asp_{sin}$, $asp_{cos}$, bs = "cc") | 0.57 | 0.37 | 1.14 | 0.91 |
| $T_{mean}$ ~ s(DOY, bs = "cc") + DEM + $asp_{sin}$ + $asp_{cos}$ + $asp_{sin}$:$asp_{cos}$ + ti(DOY, by = $asp_{sin}$, $asp_{cos}$, bs = "cc") | 0.49 | 0.27 | 1.25 | 1.00 |
| $T_{mean}$ ~ s(DOY, bs = "cc") + $asp_{sin}$ + $asp_{cos}$ + $asp_{sin}$:$asp_{cos}$ | 0.01 | 0.08 | 1.69 | 1.47 |
| $T_{mean}$ ~ s(DOY, bs = "cc") + $asp_{sin}$ + $asp_{cos}$ + $asp_{sin}$:$asp_{cos}$ + ti(DOY, by = $asp_{sin}$, $asp_{cos}$, bs = "cc") | 0.01 | 0.10 | 1.71 | 1.49 |
| **Rain** | | | | |
| **Model formula** | **Moran's I (*p*)** | **Geary's C (*p*)** | **RMSE** | **MAE** |
| $Rain_{sum}$ ~ s(DOY, bs = "cc") + DEM + ti(DOY, by = DEM, bs = "cc") | 0.82 | 0.68 | 5.10 | 2.29 |
| $Rain_{sum}$ ~ s(DOY, bs = "cc") + DEM | 0.81 | 0.59 | 5.10 | 2.32 |
| $Rain_{sum}$ ~ s(DOY, bs = "cc") + DEM + ti(DOY, by = DEM, bs = "cc") + $asp_{sin}$ + $asp_{cos}$ + $asp_{sin}$: $asp_{cos}$ | 0.91 | 0.87 | 5.18 | 2.29 |
| $Rain_{sum}$ ~ s(DOY, bs = "cc") + DEM + ti(DOY, by = DEM, bs = "cc") + $asp_{sin}$ + $asp_{cos}$ + $asp_{sin}$: $asp_{cos}$ + ti(DOY, by = $asp_{sin}$, $asp_{cos}$, bs = "cc") | 0.89 | 0.83 | 5.24 | 2.29 |
| $Rain_{sum}$ ~ s(DOY, bs = "cc") + DEM + $asp_{sin}$ + $asp_{cos}$ + $asp_{sin}$:$asp_{cos}$ | 0.90 | 0.81 | 5.24 | 2.32 |
| $Rain_{sum}$ ~ s(DOY, bs = "cc") + DEM + $asp_{sin}$ + $asp_{cos}$ + $asp_{sin}$:$asp_{cos}$ + ti(DOY, by = $asp_{sin}$, $asp_{cos}$, bs = "cc") | 0.88 | 0.82 | 5.45 | 2.49 |
| $Rain_{sum}$ ~ s(DOY, bs = "cc") + $asp_{sin}$ + $asp_{cos}$ + $asp_{sin}$:$asp_{cos}$ | 0.28 | 0.34 | >10^9 | >10^9 |
| $Rain_{sum}$ ~ s(DOY, bs = "cc") + $asp_{sin}$ + $asp_{cos}$ + $asp_{sin}$:$asp_{cos}$ + ti(DOY, by = $asp_{sin}$, $asp_{cos}$, bs = "cc") | 0.31 | 0.42 | >10^9 | >10^9 |
| **Relative humidity** | | | | |
| **Model formula** | **Moran's I (*p*)** | **Geary's C (*p*)** | **RMSE** | **MAE** |
| RH ~ s(DOY, bs = "cc") + DEM + ti(DOY, by = DEM, bs = "cc") | 0.03 | 0.05 | 0.08 | 0.06 |
| RH ~ s(DOY, bs = "cc") + DEM + ti(DOY, by = DEM, bs = "cc") + $asp_{sin}$ + $asp_{cos}$ + $asp_{sin}$: $asp_{cos}$ | 0.61 | 0.62 | 0.08 | 0.06 |
| RH ~ s(DOY, bs = "cc") + DEM | 0.02 | 0.03 | 0.08 | 0.06 |
| RH ~ s(DOY, bs = "cc") + DEM + $asp_{sin}$ + $asp_{cos}$ + $asp_{sin}$: $asp_{cos}$ | 0.53 | 0.50 | 0.08 | 0.06 |
| RH ~ s(DOY, bs = "cc") + DEM + $asp_{sin}$ + $asp_{cos}$ + $asp_{sin}$:$asp_{cos}$ + ti(DOY, by = $asp_{sin}$, $asp_{cos}$, bs = "cc") | 0.63 | 0.73 | 0.08 | 0.06 |
| RH ~ s(DOY, bs = "cc") + DEM + ti(DOY, by = DEM, bs = "cc") + $asp_{sin}$ + $asp_{cos}$ + $asp_{sin}$:$asp_{cos}$ + ti(DOY, by = $asp_{sin}$, $asp_{cos}$, bs = "cc") | 0.61 | 0.75 | 0.09 | 0.07 |
| RH ~ s(DOY, bs = "cc") + $asp_{sin}$ + $asp_{cos}$ + $asp_{sin}$:$asp_{cos}$ | 0.18 | 0.07 | 0.09 | 0.08 |
| RH ~ s(DOY, bs = "cc") + $asp_{sin}$ + $asp_{cos}$ + $asp_{sin}$:$asp_{cos}$ + ti(DOY, by = $asp_{sin}$, $asp_{cos}$, bs = "cc") | 0.31 | 0.19 | 0.10 | 0.08 |
| **Solar radiation** | | | | |
| **Model formula** | **Moran's I (*p*)** | **Geary's C (*p*)** | **RMSE** | **MAE** |
| $Rad_{mean}$ ~ s(DOY, bs = "cc") + DEM + ti(DOY, by = DEM, bs = "cc") | 0.63 | 0.56 | 62.97 | 51.81 |
| $Rad_{mean}$ ~ s(DOY, bs = "cc") + DEM + ti(DOY, by = DEM, bs = "cc") + $asp_{sin}$ + $asp_{cos}$ + $asp_{sin}$:$asp_{cos}$ | 0.76 | 0.66 | 63.67 | 52.07 |
| $Rad_{mean}$ ~ s(DOY, bs = "cc") + DEM | 0.66 | 0.59 | 64.33 | 52.88 |
| $Rad_{mean}$ ~ s(DOY, bs = "cc") + DEM + ti(DOY, by = DEM, bs = "cc") + $asp_{sin}$ + $asp_{cos}$ + $asp_{sin}$:$asp_{cos}$ + ti(DOY, by = $asp_{sin}$, $asp_{cos}$, bs = "cc") | 0.75 | 0.65 | 64.38 | 52.54 |
| $Rad_{mean}$ ~ s(DOY, bs = "cc") + DEM + $asp_{sin}$ + $asp_{cos}$ + $asp_{sin}$:$asp_{cos}$ | 0.79 | 0.70 | 64.68 | 52.90 |
| $Rad_{mean}$ ~ s(DOY, bs = "cc") + DEM + $asp_{sin}$ + $asp_{cos}$ + $asp_{sin}$:$asp_{cos}$ + ti(DOY, by = $asp_{sin}$, $asp_{cos}$, bs = "cc") | 0.72 | 0.60 | 66.75 | 54.58 |
| $Rad_{mean}$ ~ s(DOY, bs = "cc") + $asp_{sin}$ + $asp_{cos}$ + $asp_{sin}$:$asp_{cos}$ | 0.01 | 0.03 | 83.69 | 70.80 |
| $Rad_{mean}$ ~ s(DOY, bs = "cc") + $asp_{sin}$ + $asp_{cos}$ + $asp_{sin}$:$asp_{cos}$ + ti(DOY, by = $asp_{sin}$, $asp_{cos}$, bs = "cc") | 0.02 | 0.03 | 86.06 | 72.49 |

**Appendix S8. Akaike Information Criterion (AIC) values for candidate movement models.** Comparison of constant ($AIC_{const}$), single sigmoid ($AIC_{single}$), and double sigmoid ($AIC_{double}$) models fitted to individual altitudinal trajectories. The model with the lowest AIC value indicates the best-supported strategy for each individual. "No convergence" indicates models that failed to converge.

| ID | $AIC_{const}$ | $AIC_{single}$ | $AIC_{double}$ |
|---|---|---|---|
| WSN_18 | 321.37 | 325.50 | 304.59 |
| WSN_23 | 204.71 | 192.31 | 198.76 |
| WSN_28 | 260.63 | No convergence | No convergence |
| WSN_29 | 257.18 | 237.73 | 247.35 |
| WSN_31 | 360.05 | 349.87 | 321.37 |
| WSN_33 | 249.18 | 239.09 | 194.15 |
| WSN_36 | 645.18 | 613.77 | 449.25 |
| WSN_38 | 268.21 | No convergence | No convergence |
| WSN_39 | 460.49 | 460.86 | 360.41 |
| WSN_40 | 342.65 | 339.98 | 308.55 |
| WSN_41 | 144.83 | No convergence | No convergence |
| WSN_43 | 269.14 | No convergence | No convergence |
| WSN_46 | 229.54 | 219.22 | 196.13 |
| WSN_48 | 645.30 | No convergence | No convergence |
| WSN_51 | 500.02 | 372.52 | No convergence |
| WSN_53 | 326.64 | 324.47 | 257.57 |
| WSN_54 | 183.44 | No convergence | No convergence |
| WSN_70 | 664.23 | 503.33 | 603.03 |
| WSN_74 | 700.50 | 588.41 | 529.18 |
| WSN_75 | 580.87 | 432.97 | 471.75 |
| WSN_81 | 490.37 | 323.54 | 421.31 |
| WSN_84 | 462.23 | 384.39 | 398.90 |

**Appendix S9. Individual altitudinal trajectories and movement model fits for tracked *C. marthae* individuals.** Each panel displays the elevation profile (m asl) of a single individual across the annual cycle (day of year, DOY). Points represent observed GPS data, while solid lines indicate the best-fitting movement model (stationary, single-sigmoid, or double-sigmoid). Red shaded areas represent the estimated temporal windows for migration events (Est. migration) derived from model parameters. Color-coding distinguishes between males and females.

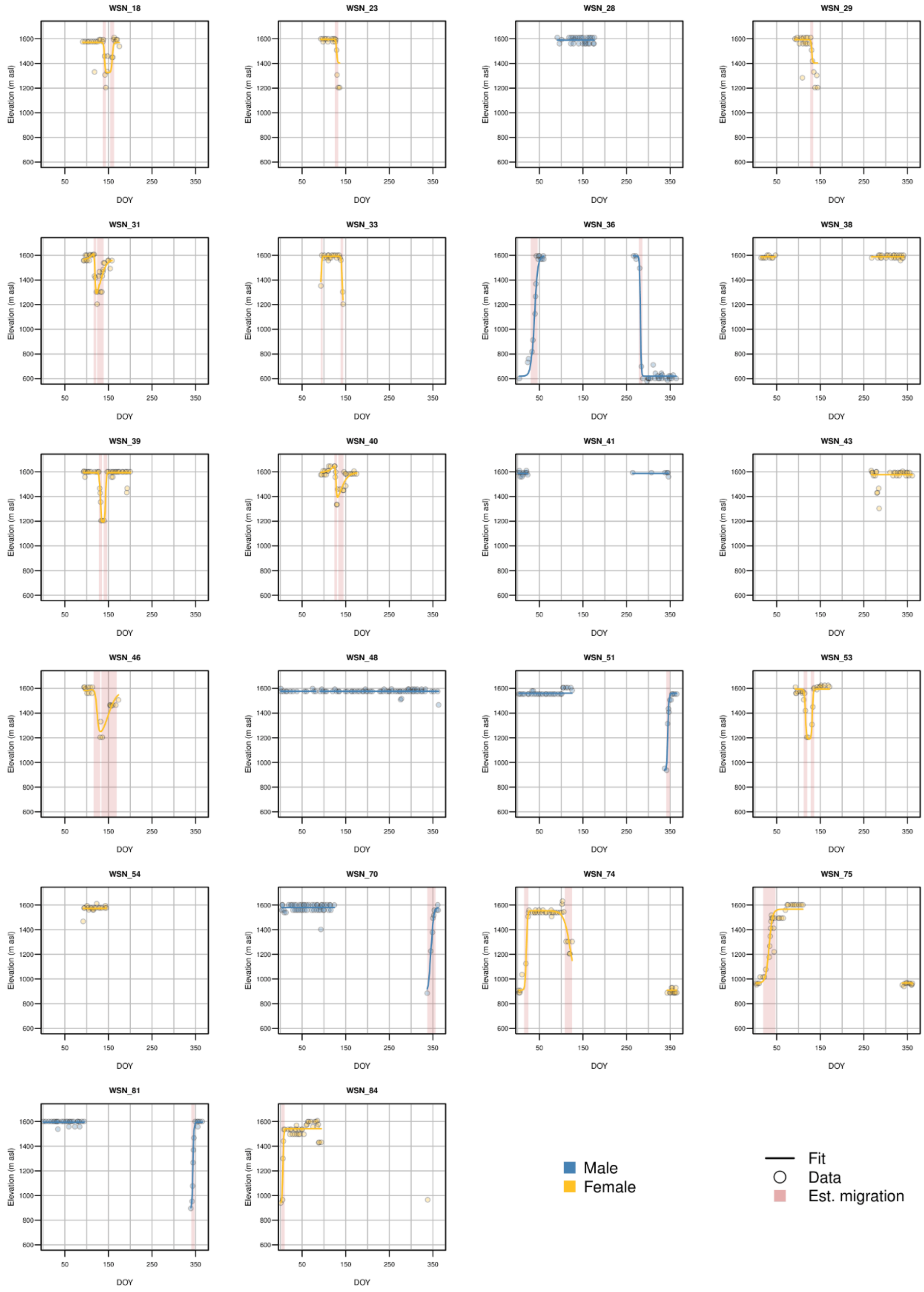

**Appendix S10. Phenological parameters and altitudinal displacement estimated for tracked *C. marthae* individuals.** For each individual (ID), the table reports the best-fitting movement model: stationary behaviour (const), monotonic (single), or bidirectional (double) movements. The estimated timing parameters for mating (Mat), nesting (Nest), and dispersal (Disp) phases are expressed in day of year (Julian days). Altitudinal shifts are categorized into seasonal migrations between mating and dispersal areas ($Shift_{MIG}$) and nesting-related movements ($Shift_{NEST}$), expressed in meters. For single-sigmoid models, the sign of the altitudinal shift indicates the direction of movement (positive for upward, negative for downward).

| ID | Sex | Model | $Mat_{START}$ | $Mat_{END}$ | $Nest_{START}$ | $Nest_{END}$ | $Disp_{START}$ | $Disp_{END}$ | $Shift_{MIG}$ | $Shift_{NEST}$ |
|---|---|---|---|---|---|---|---|---|---|---|
| WSN_18 | F | double | / | 137 | 144 | 154 | / | / | / | 244,5 |
| WSN_23 | F | single | / | 125 | 133 | / | / | / | / | -190,2 |
| WSN_28 | M | const | / | / | / | / | / | / | / | / |
| WSN_29 | F | single | / | 127 | 134 | / | / | / | / | -187,9 |
| WSN_31 | F | double | / | 116 | 122 | 124 | / | / | / | 403,8 |
| WSN_33 | F | double | 97 | 138 | 144 | / | / | / | / | 388 |
| WSN_36 | M | double | 46 | / | / | / | 286 | 30 | 975 | / |
| WSN_38 | F | const | / | / | / | / | / | / | / | / |
| WSN_39 | F | double | / | 128 | 135 | 139 | / | / | / | 402 |
| WSN_40 | F | double | / | 124 | 130 | 133 | / | / | / | 311,4 |
| WSN_41 | M | const | / | / | / | / | / | / | / | / |
| WSN_43 | F | const | / | / | / | / | / | / | / | / |
| WSN_46 | F | double | / | 116 | 131 | 134 | / | / | / | 406,5 |
| WSN_48 | M | const | / | / | / | / | / | / | / | / |
| WSN_51 | M | single | 349 | / | / | / | / | 341 | 625,2 | / |
| WSN_53 | F | double | / | 112 | 120 | 128 | / | / | / | 406,1 |
| WSN_54 | F | const | / | / | / | / | / | / | / | / |
| WSN_70 | M | single | 356 | / | / | / | / | 337 | 694,7 | / |
| WSN_74 | F | double | 25 | 108 | 125 | / | / | 15 | 644,2 | / |
| WSN_75 | F | single | 47 | / | / | / | / | 19 | 598,8 | / |
| WSN_81 | M | single | 348 | / | / | / | / | 340 | 698 | / |
| WSN_84 | F | single | 10 | / | / | / | / | 2 | 602,7 | / |

**Appendix S11. Descriptive metrics used to quantify spatial aggregation of capture locations during mating and dispersal seasons.** For each season, the table reports the number of locations (*n*), the area of the common spatial reference window (Ref. area), the observed mean nearest-neighbour distance (Obs. NN), the expected mean nearest-neighbour distance under complete spatial randomness (Exp. NN), the resulting nearest neighbour index (NNI), and the 100% minimum convex polygon (MCP) area. Lower NNI values indicate stronger spatial aggregation.

| **Season** | ***n*** | **Ref. area ($km^2$)** | **Obs. NN (m)** | **Exp. NN (m)** | **NNI** | **MCP ($m^2$)** |
|---|---|---|---|---|---|---|
| mating | 38 | 33.49 | 63.77 | 469.39 | 0.136 | 156.32 |
| dispersal | 38 | 33.49 | 131.29 | 469.39 | 0.280 | 984.70 |

**Appendix S12. Results of linear models testing differences in environmental conditions between functional areas, seasons, and sexes in *C. marthae*.** The table reports the model summaries for NDVI and air temperature (scaled values), including estimated coefficients, standard errors (SE), t values, and associated p-values. Models included area (mating vs dispersal), season (mating vs dispersal season), sex, and their interactions. Interaction terms retained in the final models were selected based on likelihood ratio tests using a stepwise model comparison procedure.

| **NDVI** | | | | |
|---|---|---|---|---|
| **Terms** | **Estimate** | **SE** | **T value** | **p-value** |
| **Intercept** | 0.370 | 0.033 | 11.350 | < 0.001 |
| **sex(M)** | -0.375 | 0.038 | -9.962 | < 0.001 |
| **season(dispersal)** | -1.712 | 0.041 | -41.897 | < 0.001 |
| **area(dispersal)** | 0.963 | 0.029 | 33.745 | < 0.001 |
| **sex(M):season(dispersal)** | 0.146 | 0.060 | 2.428 | 0.015 |
| **Temperature** | | | | |
| **Terms** | **Estimate** | **SE** | **T value** | **p-value** |
| **Intercept** | -0.790 | 0.034 | -23.570 | < 0.001 |
| **sex(M)** | -0.185 | 0.041 | -4.501 | < 0.001 |
| **season(dispersal)** | -0.186 | 0.044 | -4.267 | < 0.001 |
| **area(dispersal)** | 2.239 | 0.044 | 50.816 | < 0.001 |
| **sex(M):season(dispersal)** | 0.511 | 0.052 | 9.878 | < 0.001 |
| **sex(M):area(dispersal)** | -0.187 | 0.050 | -3.712 | < 0.001 |
| **season(dispersal):area(dispersal)** | -0.944 | 0.052 | -18.311 | < 0.001 |
